# Deep Learning Based Cell Segmentation of 3D Confocal Images of Wood for Enzymatic Hydrolysis Applications

**DOI:** 10.64898/2026.10.26.761974

**Authors:** Ali Faraj, Solmaz Hossein Khani, Gabriel Paës, Yassin Refahi

## Abstract

Plant cell walls are an abundant renewable resource for biobased products, but their recalcitrance limits enzymatic hydrolysis efficiency. Cell and tissue morphological parameters provide reliable biomarkers for predicting enzymatic hydrolysis efficiency. Accurate cell segmentation in 3D microscopy images is essential for quantifying these parameters. Although deep learning has achieved high performance in biological image segmentation, confocal microscopy’s limited penetration depth leaves elongated wood cells partially imaged, creating open-ended structures that challenge existing segmentation methods. Based on an artificial longitudinal domain restriction, an adapted framework is presented here for the training of a 3D U-net model for the cell segmentation of wood images. Quantitative evaluation on a dedicated benchmark dataset identified a model configuration that reduced segmentation errors and produced more robust predictions during early enzymatic hydrolysis. These results support the use of the framework to quantify morphological parameters relevant to assessing and predicting biomass conversion efficiency.

## 1. INTRODUCTION

Plant cell walls are an abundant renewable resource for producing biobased products. Biotechnological conversion of cell walls relies on enzymatic hydrolysis to release simple sugars from polysaccharides. However, hydrolysis efficiency remains limited by cell wall recalcitrance. Therefore, it is essential to better understand enzymatic hydrolysis to achieve a better conversion efficiency. Cell and tissue scales parameters have been shown to predict hydrolysis yields, showing their potential as indicators of conversion efficiency. Despite this potential, enzymatic deconstruction at the micro scale remains under-investigated compared to the nanoscale. This gap is largely due to the challenges faced in analyzing cell wall images during hydrolysis. Confocal laser scanning microscopy (CLSM) makes it possible to acquire three-dimensional imaging of cell walls during deconstruction. Segmentation allows identification of individual cells from the collected images.

Existing analysis pipelines propagate spatial information from a segmented reference image acquired at an earlier time-point to later hydrolysis-phase images [1, 2]. The segmentation accuracy of these later images therefore depends on the quality of the reference segmentation. In these pipelines, initial segmentation errors are propagated through the time-lapse images and may lead to significant errors in subsequent hydrolysis-phase segmentations. Obtaining a reliable reference segmentation requires time-consuming manual corrections, making these pipelines difficult to apply to large datasets. Accurate automated 3D cell segmentation is thus needed to reduce manual intervention.

Recent deep learning methods achieve high segmentation accuracy while reducing manual intervention after training. Deep learning models based on Convolutional Neural Networks (CNN) have shown their efficiency as boundary predictors. This was used for the development of the U-Net architecture for the segmentation of 2D biomedical images [3] and was extended to segment 3D images [4]. The combination of U-Net architectures with classical (non-deep learning) algorithms has shown a strong potential for the image segmentation of cells [5, 6]. In particular, thanks to a Pytorch implementation of the 3D U-Net architecture, the authors in [5] could perform the cell segmentation of various plant tissues and organs with an excellent performance. This Pytorch 3D U-Net could provide more accurate and more robust results compared to other deep learning and classical algorithms for 3D cell segmentation [7]. For this reason, we propose a training strategy of the Pytorch 3D U-Net model for the cell segmentation of wood confocal images at a fixed time of the enzymatic hydrolysis.

Confocal microscopy enables high-quality 3D reconstruction through precise optical sectioning and resolution down to ≈ 0.2 *µm*, but its limited penetration depth (typically ≤ 50 *µm*) results in only partial imaging of elongated wood cells, creating open-ended structures (Fig. 2.A), which challenges the direct application of the existing deep learning-based segmentation methods. The training strategy presented here is based on a data preparation suited for this specific cell-shape. This is obtained by artificially closing the imaged samples in the longitudinal direction to avoid cell merge events due to the presence of open-cells in this direction.

**Fig 1.**
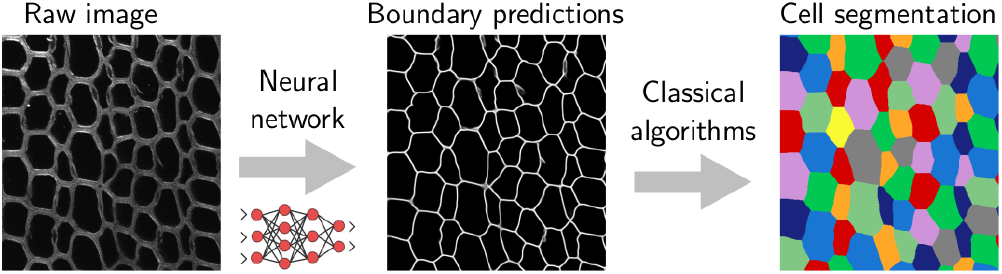
Representation of the Unet+watershed and PlantSeg pipelines. In the first step, both pipelines use the same 3D U-Net to predict the cell boundaries from the raw images. In the second step the pipelines use classical algorithms (watershed for Unet+watershed and volume partitioning for PlantSeg) to segment the cells from the boundary predictions.

**Fig 2.**
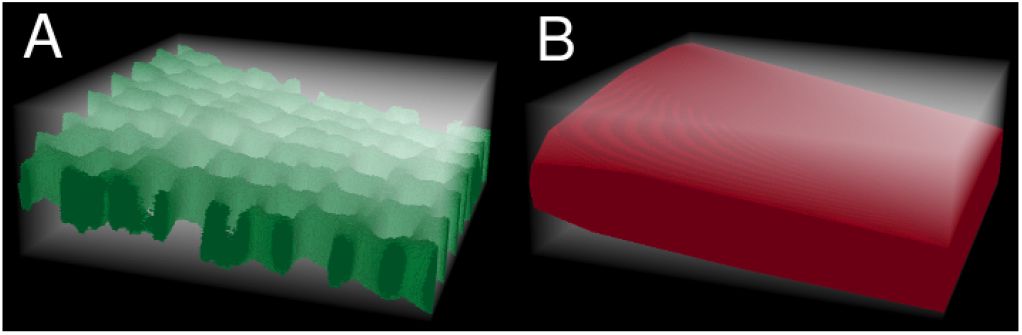
Pre-hydrolysis 3D acquisition of a tilted wood sample. (A) Raw image of the wood sample exhibiting open-ended cells. (B) 3D convex hull of the wood sample depicted in red. In A–B, the semi-transparent white color represents the background.

Since the watershed algorithm proved to perform well for instance segmentation, watershed techniques were combined with different deep learning models, including 3D U-Net [8]. However, existing pipelines are not always available, compromising model re-training on new data. In the present work, the deep learning based segmentation employs the Pytorch 3D U-Net model to predict cell boundaries and the watershed algorithm to deduce instance cell segmentation from the predicted boundaries. The resulting pipeline was bench-marked against the PlantSeg pipeline [5] on a dedicated, expert-curated dataset of wood enzymatic hydrolysis. Metrics to evaluate the accuracy of the predicted cell boundaries and segmentation, including over- and undersegmentation rates, were implemented and adapted to the specific shape of wood cells.

In this study, we compare different training strategies for the PyTorch 3D U-Net in order to identify an efficient approach for predicting cell boundaries in 3D images of wood acquired using confocal fluorescence microscope, where the limited imaging depth results in partially imaged, open-ended cells. In particular, the prediction accuracy provided by the two following training strategies were compared: use a pretrained model from [5] that could benefit from a variety of large datasets during the training step; or train the model from scratch on our more specific wood dataset (fine-tuning is not presented here since it did not improve the prediction accuracy). For the training on our own data, the prediction accuracy related to the two following choices of the training set were compared: training on pre-hydrolysis images; or training on images from the early hydrolysis-stage. This is complemented with a visualization of segmentation errors by time points to evaluate if the training data augmentation with hydrolysis-stage information could improve the predicted segmentation at the hydrolysis-stage, without recourse to a propagation strategy of pre-hydrolysis segmentation.

## 2. DEEP LEARNING BASED SEGMENTATION AND PREPROCESSING OF WOOD IMAGES

To compute the deep learning based segmentation, two main steps were performed. In the first step, the 3D U-Net is trained for the prediction of cell boundaries in 3D (through voxel-wise boundary classification). In the second step, the predicted cell boundaries are used to compute the final instance segmentation (see Fig. 1). For the second step, the strategy proposed here is to perform a watershed algorithm on the voxel-wise probabilities provided by the 3D U-Net predictions, in order to deduce the instance cell segmentation. The watershed algorithm is preceded by a Gaussian filter in order to limit oversegmentation. The resulting pipeline will be called Unet+watershed. In [5], the second step is obtained by performing a volume partitioning algorithm to segment the cells from the boundary predictions. The resulting segmentation algorithm corresponds to the PlantSeg pipeline of [5].

Because wood cells are elongated and extend beyond the sampled section or confocal imaging depth, only portions of their length are captured, leaving cell lumens open at the boundaries of the imaged volume. As a consequence, the input images of the 3D U-Net present connected cells leading undersegmentation. To overcome this issue, the deep learning based segmentation pipelines were restricted to the region inside the convex hull of the sample. Indeed, such a restriction is more suited than a domain restriction related to given longitudinal positions because of the presence of titled or deformed samples. The sample convex hull is computed as explained in [1]. For the cell boundary prediction step (Step 1), the information outside the convex hull is not relevant and might lead to incorrect predictions. Therefore, voxels outside the sample’s convex hull were excluded before the 3D U-Net training and inference by setting their value to 0 in the raw images and to an unused label in the ground truth segmentations.

The instance segmentation step (Step 2) is computed from the 3D U-Net predicted probabilities which describe the boundaries of cells with possibly open-ends. Therefore, the classical algorithm of Step 2 (non-deep learning) should be restricted to the sample convex hull to succeed, as explained above. For this reason, the Step 2 was pre-processed by setting the voxels of the predicted probabilities to the value 1 (maximal probability) outside the convex hull. The open-ended shape of wood cells as well as the convex hull of a tilted sample are shown in Fig. 2.

## 3. DATASET

Spruce (Picea abies) wood samples were collected, prepared and imaged by following the protocol in [1, 2, 9]. Specifically, time-lapse imaging of the spruce samples during enzymatic hydrolysis was performed with the Olympus FV3000 confocal microscope generating sequences of 3D images with a voxel size of 0.62 *×* 0.62 *×* 0.3 *µm*. Each generated sequence includes 25 times points of size 512 *×* 512 *×* 400 voxels corresponding to hourly acquisitions. At late time points, individual cells cannot be isolated due to the loss of integrity of cell walls caused by the enzymatic hydrolysis and, both classical and deep learning, segmentation algorithms cannot be used directly [1, 2]. For this reason, the two datasets considered here correspond to images collected during the first 5 hours of the enzymatic hydrolysis. The pre-hydrolysis dataset (Hyd-0h) includes the initial image from each time-lapse sequence, with 26 images randomly assigned to training, 3 to validation and 6 to testing. The hydrolysis time-course dataset (Hyd-0–5h) includes images acquired from 0 to 5 h, with 153 images randomly assigned to training, 18 to validation and 33 to testing. To allow comparisons, we imposed that the Hyd-0–5h test set contains the Hyd-0h test images.

The benchmark segmentation (ground truth) [10] is computed with the classical (non-deep learning) pipeline MARS [11], combined with a manual correction by an expert of the segmentation errors, as explained in [12].

## 4. TRAINING

For the training process, the ground truth cell boundaries are obtained from the ground truth segmentation by finding two voxels-thick boundaries between labeled regions as explained in [5]. Training of the 3D U-Net was performed with the Adam optimizer with a weight decay of 0.00001 and the initial learning rate *ε* = 0.0002. Training with small patches of size 40 *×* 85 *×* 85 was selected (the same patch size was used during inference to preserve the prediction quality). The training of the model required 150 000 iterations before convergence on a NVIDIA H100 GPU and could benefit from 16 CPU cores. The model was trained using the sum of the Binary Cross-Entropy and Dice loss functions (*L* = *L*_*BCE*_ + *L*_*Dice*_). The data was augmented using random flips, random rotations, elastic deformations and noise augmentations. The evaluation metrics used for validation is the Adapted Rand Error [13].

## 5. EVALUATION METRICS

The metrics used to evaluate the accuracy of the boundary predictions are the Accuracy, Precision, Recall and F1 scores. The Accuracy denotes the percentage of correctly predicted voxels. For the final segmentation, the Volume averaged Index (*VI*) in [7] was used to quantify the degree of overlap between the ground truth segmentation *G* = {*G*_*i*_}and the predicted segmentation *P* = {*P*_*j*_} . The *VI* index is obtained by comparing each cell *G*_*i*_ in the ground truth to the cell in *P* of best overlap:

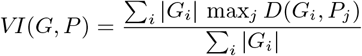

where *D*(*A, B*) is the Dice Similarity Index of the sets *A* and *B* and the sums are (resp. the max function is) taken over all the indices of *G* (resp. of *P*) except the background index. The rates of over- and undersegmentation are quantified through a cell by cell computation of the asymmetric inclusion index:

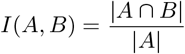

Each cell *G*_*i*_ is associated with a cell of index *j* = *B*(*i*) in *P* according to the maximal inclusion:

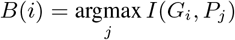

Similarly, each cell *P*_*j*_ is associated with a cell of index *i* = *B*^*′*^(*j*) in *G* where:

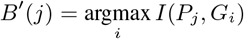

Based on the above associations, a binary graph is built which is then used to evaluate the state of each cell *P*_*j*_ :

- one-to-one: the cell *P*_*j*_ is only connected to the cell *G*_*B′*(*j*)_
- undersegmentation: *P*_*j*_ is connected to several cells of *G* which are not connected to any other cell of *P*
- oversegmentation: *G*_*B′*(*j*)_ is connected to several cells of *P* which are not connected to any other cell of *G*

The rates of over- and undersegmentation are obtained by counting the number of cells of *P* in each state. The cells of *P* which are associated with the background of *G* are not counted in the rate computation (see [7] and [14] for more details).

As a consequence of section 2, all the metrics presented here were computed by discarding the voxels outside the convex hull.

## 6. RESULTS

The accuracy of the Unet+watershed and PlantSeg pipelines was evaluated at the two steps of the pipelines. For both pipelines, the following training strategies were considered: use a pretrained model (3D-Unet-pretrained), train the model on the Hyd-0h training set (3D-Unet-Hyd-0h), and train the model on the Hyd-0–5h training set (3D-Unet-Hyd-0–5h). The pretrained model was the `generic confocal 3D unet` provided by [5].

### 6.1. Step 1: Cell boundary prediction

The boundary prediction was first evaluated on the images of the testing set of the Hyd-0h dataset (Hyd-0h testing set). Then, to evaluate the accuracy on the hydrolysis-stage images, the boundary prediction was evaluated on the subset of the test images of Hyd-0–5h corresponding to the time points t=1–5h. This subset will be denoted “Hyd-1–5h testing set” and contains 23 images. The predicted cell boundary binary masks were deduced by thresholding the probabilities output by the 3D U-Net. For each model and for each testing set, the threshold was chosen by optimizing the F1 score on the corresponding training and validation data. The resulting boundary prediction scores are presented in Table 1. It can be noticed that retraining the 3D U-Net model on our own data improved the cell boundary predictions. Whereas, increasing the pre-hydrolysis training set (Hyd-0h) with the hydrolysis-stage training set (Hyd-0–5h) did not improve the cell boundary predictions, even on the hydrolysis-stage Hyd-1–5h testing set. Moreover, the 3D-Unet-Hyd-0h model provided a higher mean Accuracy compared to the other models (96.1% on the Hyd-0h testing set and 92.2% on the Hyd-1–5h testing set). As a conclusion, the 3D-Unet-Hyd-0h model showed the best performance with accurate cell boundary predictions and metrics values in agreement with existing works [5].

**Table 1.** Boundary prediction accuracy. Mean scores over the considered testing sets are presented (a higher value is better).

| Network | Precision | Recall | F1 |
| --- | --- | --- | --- |
| Hyd-0h testing set |  |  |  |
| 3D-Unet-pretrained | 0.390 | 0.516 | 0.444 |
| 3D-Unet-Hyd-0h | 0.653 | 0.740 | 0.693 |
| 3D-Unet-Hyd-0–5h | 0.540 | 0.702 | 0.610 |
| Hyd-1–5h testing set |  |  |  |
| 3D-Unet-pretrained | 0.297 | 0.535 | 0.382 |
| 3D-Unet-Hyd-0h | 0.405 | 0.609 | 0.487 |
| 3D-Unet-Hyd-0–5h | 0.390 | 0.647 | 0.486 |

### 6.2. Step 2: Cell segmentation

As a consequence of the above results, Step 1 was performed with the 3D-Unet-Hyd-0h model for both Unet+watershed and PlantSeg (the 3D U-Nets of these two pipelines share the same coefficients). Then, the final segmentation accuracy of the Unet+watershed and PlantSeg pipelines was evaluated against the ground truth segmentation. We also investigated the impact of the tuning of Step 2 parameters on the overall segmentation performance of both pipelines, without considering a systematic parameter optimization.

For the Step 2 of the Unet+watershed pipeline, two parameters were considered for the hyperparameter tuning: the parameters *σ* of the Gaussian filter and the parameter *h*_*min*_ used to extract intensity local minima and provide seeds to the watershed algorithm. The default value of *σ* and *h*_*min*_ (denoted as Unet+Ws-default) was selected thanks to the MARS pipeline used to compute the ground truth segmentation [12]. After the hyperparameter tuning, the values *σ* = 5 and *h*_*min*_ = 10 were selected (denoted as Unet+Ws-tuned). In the PlantSeg pipeline, the Multicut algorithm was selected for Step 2. The PlantSeg segmentation depends on several parameters that are mainly related to the super-voxels generation [5]. The tuning of only two of these parameters was considered here: the CNN prediction threshold and the over/under segmentation factor. The default value of these parameters was chosen, according to the pipeline specification, as: 0.5 and 0.5 respectively (denoted as PlantSeg-default). We obtained respectively the values 0.6 and 0.4 after the hyper-parameter tuning (denoted as PlantSeg-tuned).

As it appears from the results summarized in Table 2, the over-segmentation rate with the default segmentation parameters is important, for both Unet+watershed and PlantSeg. Therefore, the parameters of Step 2 were chosen to orient the pipelines toward undersegmentation, resulting in a reduction of the oversegmentation rate by a factor 2 for PlantSeg, and a factor 3 for Unet+watershed. At the cost of a marginal decrease of the *VI* average and a reasonable increase of the undersegmentation rate, this parameter tuning leads to a significant improvement of the rate of correctly segmented cells (One-to-one). It should be noticed that, after the parameter tuning, the overall performance of the pipeline Unet+watershed is better than that of PlantSeg. Our numerical experiments also indicate that further adjustments to the Step 2 parameters of the Unet+watershed pipeline could increase the rate of correctly segmented cells and reduce oversegmentation, at the cost of a reasonable increase in undersegmentation (Table 2). However, systematic parameter optimization was beyond the scope of this section.

**Table 2.** Segmentation accuracy. One-to-one, Overseg. and Underseg. denote respectively the rate of correctly segmented cells and the rates of over- and undersegmentation, computed on the Hyd-0h testing set.

| Network | One-to-One | Overseg. | Underseg. |
| --- | --- | --- | --- |
| Unet+Ws-default | 69.7% | 25.5% | 1.14% |
| Unet+Ws-tuned | 83.9% | 8.07% | 4.25% |
| PlantSeg-default | 74.6% | 18.7% | 2.27% |
| PlantSeg-tuned | 83.0% | 9.67% | 4.32% |

Fig. 3 shows the variation of the segmentation accuracy of the Unet+watershed and PlantSeg pipelines on the Hyd-0h and Hyd-1–5h testing sets. For the PlantSeg pipeline, the model in Step 1 is trained on the Hyd-0h training set (PlantSeg-Hyd-0h). For the Unet+watershed pipeline, two strategies were compared: the model is trained on the Hyd-0h training set (Unet+watershed-Hyd-0h) and the model is trained on the Hyd-0–5h training set (Unet+watershed-Hyd-0–5h). For the three pipelines (PlantSeg-Hyd-0h, Unet+watershed-Hyd-0h and Unet+watershed-Hyd-0–5h), the parameters of Step 2 are the tuned parameters in Table 2.

**Fig 3.**
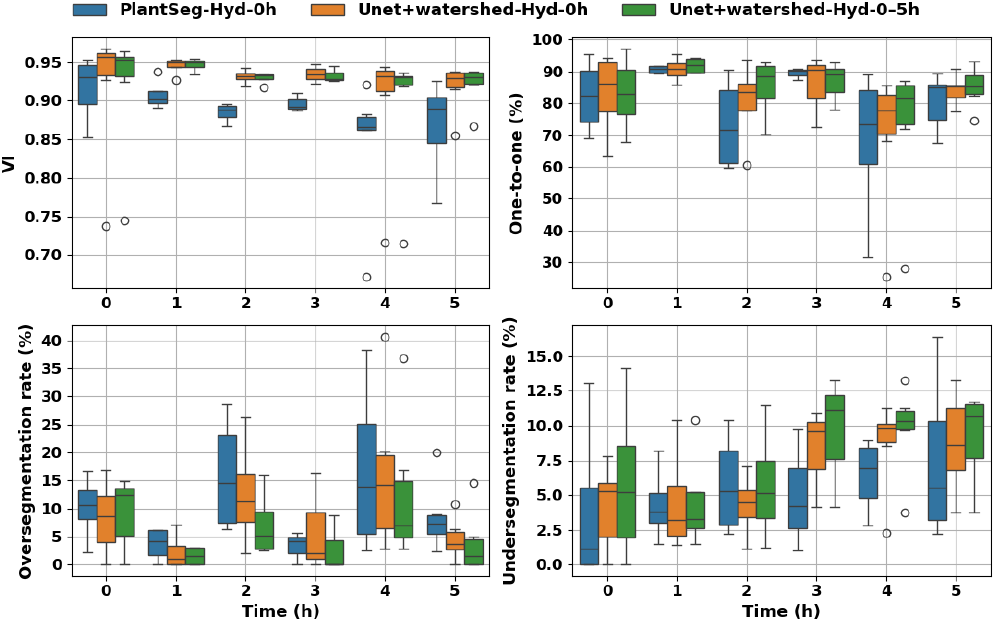
Quantification of the segmentation produced by the PlantSeg-Hyd-0h, Unet+watershed-Hyd-0h and Unet+watershed-Hyd-0–5h pipelines. The values at the time point t=0h were obtained on the Hyd-0h testing set (6 images) whereas the values at the later time points were obtained on the Hyd-1–5h testing set (4 images at t=1h, 4 images at t=2h, 3 images at t=3h, 6 images at t=4h and 6 images at t=5h).

The results in Fig. 3 confirm that, for the selected pipeline configurations, the overall accuracy of Unet+watershed outperforms that of the PlantSeg pipeline and that the increase of the training set with the hydrolysis-stage training images does not improve Unet+watershed overall accuracy. We also note that the PlantSeg pipeline has a tendency to oversegment, which is even more important for later time steps. Inversely, Unet+watershed-Hyd-0–5h tends to undersegment. This however leads to a slight improvement of the rate of correctly segmented cells for latter times (but not of the *VI* index). This tendency to undersegment can be explained by the results of Table 1. Indeed, increasing the training set with the Hyd-0–5h training images provides a lower accuracy of the 3D U-Net predictions (as already said above), leading to the possible apparition of holes in the predicted cell boundaries and, therefore, to cell merge events.

## 7. CONCLUSION

We proposed a deep learning based segmentation pipeline dedicated to the cell segmentation of wood cell images. The pipeline allows to take advantage of the cell boundaries predicted by a 3D U-Net thanks to an artificial domain restriction in the longitudinal direction, where wood cells have open-ends. Metrics to evaluate the accuracy of the predicted cell boundaries and segmentation, including over- and undersegmentation rates, were adapted to this specific cell-shape. The present work provides a training and hyperparameter tuning strategy suited to cell wood images. Specifically, re-training the deep learning part of the pipeline on our own data led to a significant improvement of the prediction accuracy compared to a pre-trained model. The use of the watershed algorithm for the deduction of the instance cell segmentation has resulted in more accurate segmentations compared to the volume partitioning algorithm in PlantSeg. After these improvements, our pipeline could provide accurate cell segmentation of pre-hydrolysis images. However, our results show that the increase of the training set with hydrolysis-stage images does not improve the predicted segmentation overall accuracy, even on hydrolysis-stage images.

To overcome this, the accurate segmentation results provided by our pipeline on pre-hydrolysis images can be used to initialize a (possibly deep learning based) cell tracking algorithm suitable for high deconstruction. Morphological similarities between spruce and other wood species, such as poplar, suggest that the predicted wood cell segmentation could benefit of a further improvement through retraining on datasets from these species. By addressing the challenges posed by partially imaged, open-ended cells, the proposed framework enables accurate 3D segmentation before and during early enzymatic hydrolysis, providing a foundation for quantifying morphological biomarkers to assess and predict biomass conversion efficiency.

## 8. COMPLIANCE WITH ETHICAL STANDARDS

This research study was conducted using plant data study for which no ethical approval was required.

## 9. ACKNOWLEDGMENTS

This work was supported by the French Ministry of Research via the Agence Nationale de la Recherche through the BIOMOD grant to Y.R. (ANR-19-CE43-0010) and through the FillingGaps grant to G.P. (ANR-23-PEBB-0006). Additional support was provided by the INRAE TRANSFORM division (DeepAnaDec to A.F.). This work could benefit from the computing resources of the ROMEO Regional Computing Center (https://romeo.univ-reims.fr). The authors thank Grégoire Malandain for his comments on the manuscript.

